# SynPull: a novel method for studying neurodegeneration-related aggregates in synaptosomes using super-resolution microscopy

**DOI:** 10.1101/2024.08.24.609517

**Authors:** Shekhar Kedia, Emre Fertan, Yunzhao Wu, Yu P. Zhang, Georg Meisl, Jeff Y. L. Lam, Francis Wiseman, William A. McEwan, Annelies Quaegebeur, Maria Grazia Spillantini, John S. H. Danial, David Klenerman

## Abstract

Synaptic dysfunction is one of the primary hallmarks of both Alzheimer’s and Parkinson’s disease, leading to cognitive and behavioural decline. While alpha-synuclein, beta-amyloid, and tau are involved in the physiological functioning of synapses, their pathological aggregation has been linked to synaptic dysfunction. However, the methodology for studying the small (sub-diffraction limit) and soluble aggregates -often called oligomers, formed by these proteins is limited. Here we describe SynPull, a novel method combining single-molecule pulldown, super-resolution microscopy, and advanced computational analyses, in order to reliably study the quantity and morphology of the oligomeric alpha-synuclein, beta-amyloid, and AT8-positive tau aggregates in synaptosomes harvested from post-mortem human brain samples and mouse models. Using SynPull, we show that AT8-positive tau is the predominant aggregate type in AD, with significantly more aggregates compared to the control samples, yet the aggregate size does not differ between disease and control samples. Meanwhile, the relatively smaller amount of alpha-synuclein and beta-amyloid aggregates found in the synapses are larger than the extra-synaptic ones. Collectively, these results show the utility of SynPull to study pathological aggregates in dementia, which can help further understand the disease mechanisms causing synaptic dysfunction.

Graphical abstract.
Human post-mortem orbitofrontal cortex samples from subjects with neuropathological diagnosis of Alzheimer’s and Parkinson’s disease, as well as age-matched controls cut into ∼300 mg sections, and MI2, APP^NL-G-F^, P301S, and C57Bl/6J mouse brains were first homogenised in synaptosome buffer using a Dounce homogeniser and then filtered and centrifuged to separate nuclei and organelles from the synaptic fragments. Then, the isolated synaptosomes were incubated on the SiMPull surface with anti-neurexin antibody overnight, followed by fixation and permeabilisation. Imaging antibodies against beta-amyloid, alpha-synuclein, and AT8-positive tau were added to the samples and *d*STORM imaging was performed to super-resolve the aggregates.

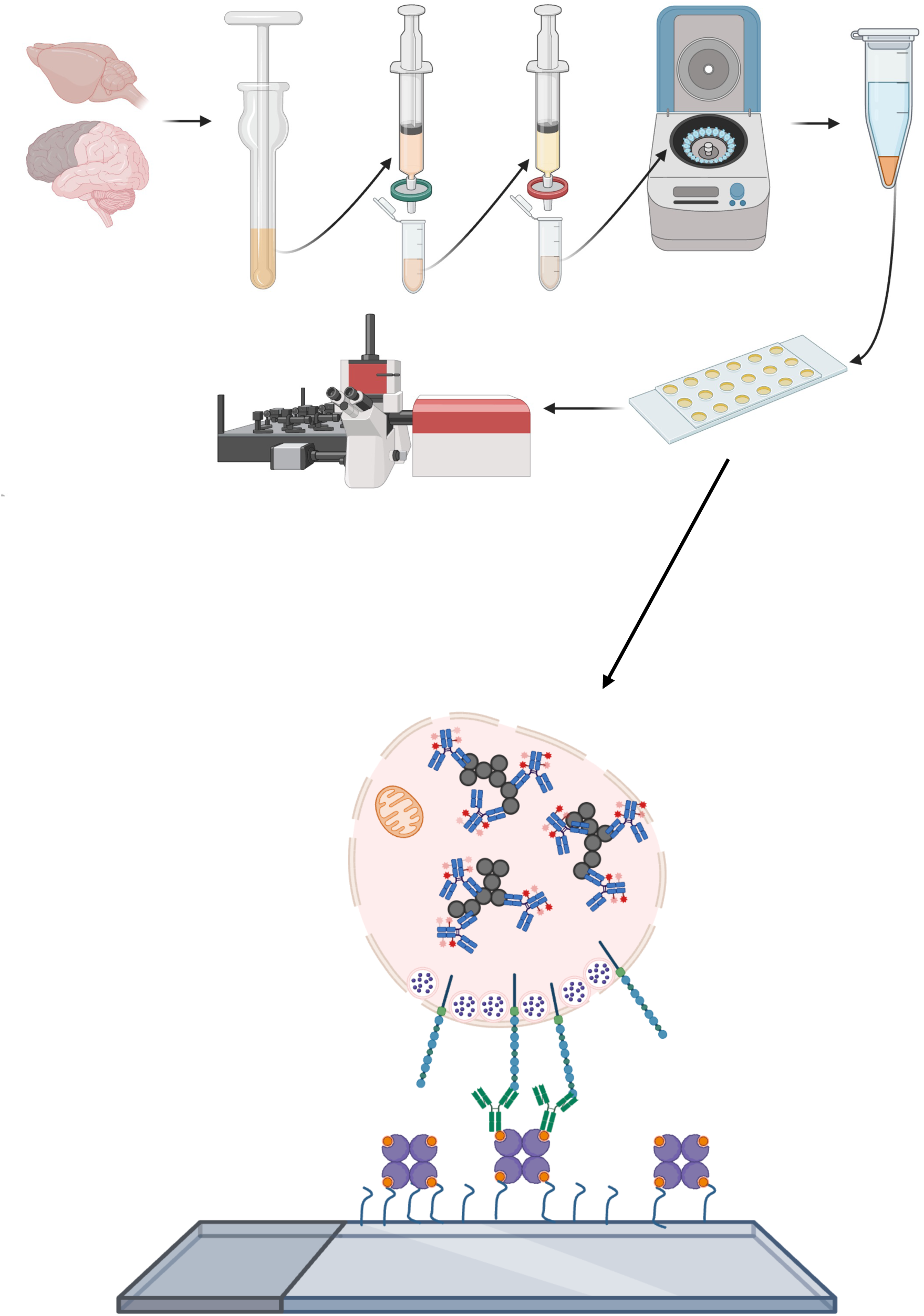

## 1. Introduction

Alzheimer’s and Parkinson’s disease (AD and PD) collectively make-up more than half of all neurodegenerative disease cases. Both are neuropathologically characterised by increased inflammation, oxidative stress, and the abnormal accumulation and aggregation of beta-amyloid (Aβ) and hyperphosphorylated (p)tau^1,2^ in AD, and alpha-synuclein (ɑSyn)^3,4^ in PD. Another common pathology seen in both diseases is progressive synaptic dysfunction, leading to cognitive and behavioural deficits. Indeed, in both diseases pathological aggregates of the proteins mentioned above have been linked to synaptic dysfunction^5–7^.

The presence and role of Aβ in the synapse is an area of active research, with some evidence suggesting a role in neuronal function, especially in times of increased synaptic activity, when Aβ production is increased^8^. When neurons were treated with β- and γ-secretase inhibitors to stop amyloid precursor protein (APP) processing through the amyloidogenic pathway, increased neuronal loss was observed, which could be ameliorated by Aβ40 treatment^9^. Soluble N-ethylmaleimide sensitive factor attachment receptors (SNARE) complexes -which are involved in synaptic neurotransmitter release, are reduced in post-mortem AD brains^10^ as well as mouse models^11^, indicating synaptic dysfunction in AD. Interestingly, the total amount of synaptobrevin, syntaxin-1, or SNAP-25, which are components of the SNARE complex, were not altered, suggesting that Aβ inhibits the formation of the SNARE complex. Indeed, Yang *et al.*^11^ showed direct binding of oligomeric Aβ to syntaxin-1, altering SNARE complex formation. The presence of Aβ is not only limited to the pre-synapse, as it has been shown that Aβ can interact with post-synaptic neuroligin (NLGN)1, leading to Aβ oligomer formation^12^. Thus, while Aβ may be a physiological component of all synapses, formation of pathological Aβ aggregates with different sizes and structures may shift its role from physiological to pathological and promote synaptic loss. Therefore, quantifying, and characterising Aβ aggregates in the synapses will be a valuable method for identifying novel pathological AD-related mechanisms.

Similarly, the presence of pTau in the synapse has also been studied. Frandemiche *et al*.^13^ showed tau is localized in the axonal dendritic shafts in healthy neurons during resting conditions and synaptic activity relocated tau to the synaptic terminals, interacting with actin. However, in the presence of oligomeric Aβ, tau was translocated to the synapse in the resting state and both tau and actin levels were decreased after synaptic activity, suggesting an Aβ-dependent synaptic dysfunction promoted by tau. Moreover, it has also been suggested that tau can interfere with synaptic vesicles. AT8-positive tau can bind to synaptic vesicles through its N-terminal domain and interfere with presynaptic functions, including synaptic vesicle mobility and release ^14^. Moreover, tau can spread between neurons via synaptic transmission^15^. and synaptic accumulation of oligomeric tau, in the absence of fibrillar species has been reported was observed in early AD, suggesting the involvement of these small-soluble aggregates in synaptotoxic processes^16^.

Lastly, ɑSyn is a synaptic protein required for synaptic function and neurotransmitter release, through its involvement in the SNARE complex formation by interacting with synaptobrevin^17^. However when ɑSyn starts to aggregate, the formed oligomers recruit the synaptic complex proteins^18^ and inhibit neurotransmitter vesicle docking and release^19,20^. While the Lewy bodies are the most recognised form of ɑSyn accumulation in PD and other synucleinopathies, recent studies have shown that the earlier forms of soluble ɑSyn aggregates, may be more toxic than the Lewy bodies^21^. Our group has recently shown that the size and shape of the ɑSyn aggregates, most of which are below the diffraction-limit of light, differs as the disease progresses and influences their toxic properties^22^ making it essential to study ɑSyn aggregates in the synapse.

As explained above, pathologically-aggregated Aβ, pTau, and ɑSyn are all associated with synaptic dysfunction. The smaller, oligomeric aggregates formed by these proteins, rather than the larger and insoluble plaques, tangles, and Lewy bodies may be the highly toxic forms, promoting synaptic loss. The sub-diffraction limit size, low abundance in the synapse, and high heterogeneity of these aggregates makes it difficult to study and characterise these small-soluble aggregates^23^ using traditional methods such as immunohistochemistry. Along with electrophysiology, one of the most valuable tools for studying synaptic pathology is synapto(neuro)somes^24^. First used by Catherine Hebb and Victor Whittaker^25^, synaptosomes are spherical membrane bound structures with a diameter ranging from 0.6 to 2 µm, made of nerve endings torn from the axon containing the synaptic boutons with neurotransmitter vesicles, synaptic proteins, mitochondria, and fragments of post-synaptic density. Here, we developed and validated SynPull, as a novel method of studying synaptosomes with single-molecule pulldown (SiMPull) and super-resolution microscopy, enabling the specific characterisation of Aβ, tau, and ɑSyn aggregates in terms of number, size, and shape from post-mortem human brain samples as well as mouse models of AD, tauopathy, and PD (**Graphical abstract**). By applying single-molecule detection techniques to aggregates located inside synaptosomes, SynPull opens up novel research avenues to study the role of pathological aggregates in synaptic dysfunction.

## 2. Results

### Electron microscopy studies

To confirm the integrity of isolated synapto(neuro)somes, electron microscopy (EM) was performed on synaptosome preparations from mouse and human brains. As shown in **Figure 1**, synaptoneurosomes containing post-synaptic compartments and synaptosomes with pre-synaptic boutons were successfully prepared from the human cortex (**Figure 1A**) and C57Bl/6J mouse cerebrum (**Figure 1B**). Morphological features such as synaptic vesicles, synaptic membranes, and mitochondria observed in these micrographs validate the integrity of the isolated synapto(neuro)somes.

**Figure 1.**
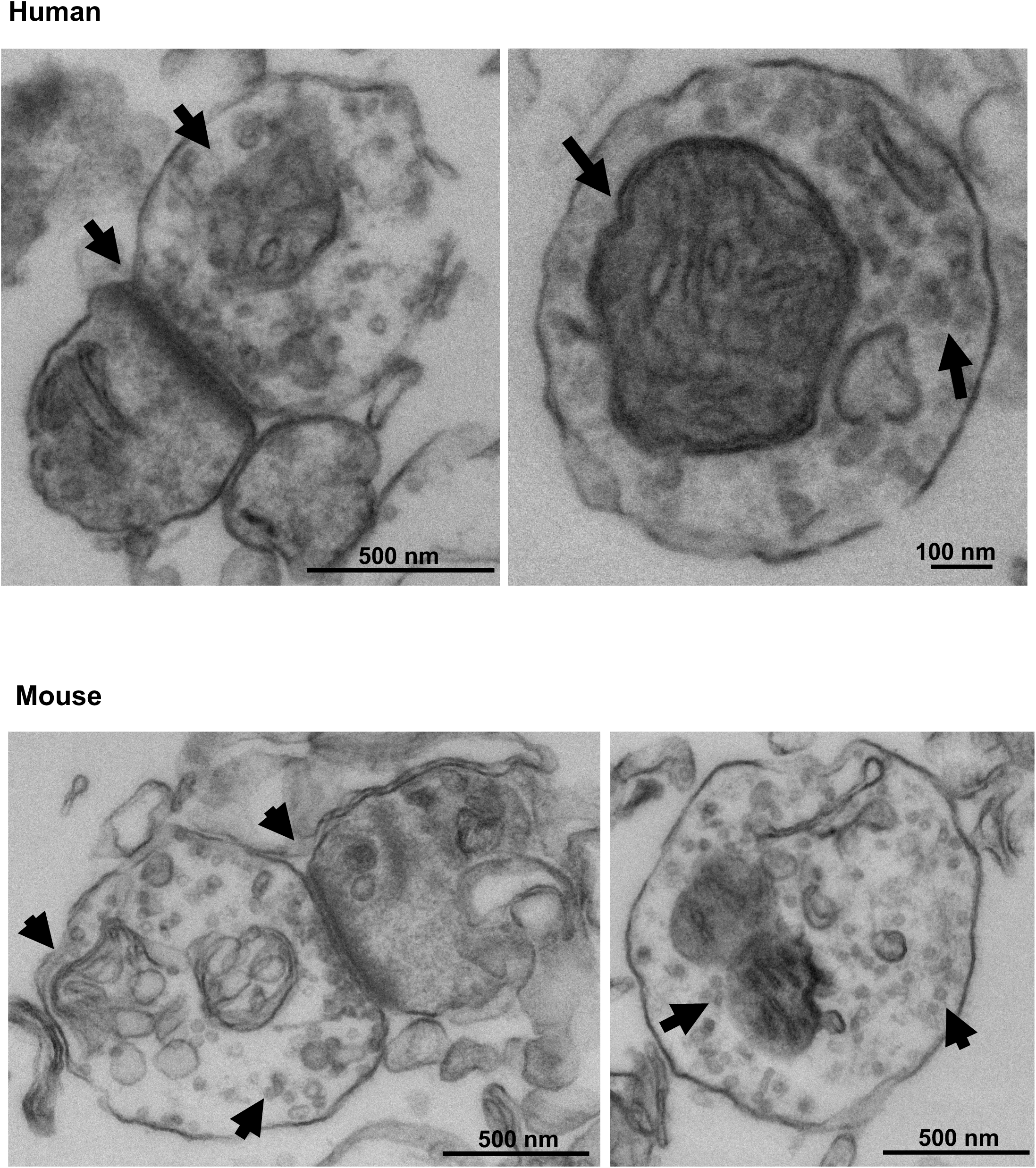
Electron micrographs of human (control) and mouse (C57Bl/6J) synaptosomes.

### Control studies

In order to validate SynPull and test the suitability of the SiMPull surface for our experiments, we performed a number of control experiments (**Figure 2A**). Since the 561 nm (green) laser is used to excite the CellMask, which is used to identify synaptosomes, it was crucial that the surface is not auto-fluorescent under illumination at this wavelength. When the surface was treated with all the steps required for the experiment, including neutravidin, polymer, NRXN1 capture antibody, fixing, and permeabilising, but no sample and CellMask added, no signal was detected under illumination with the 561 nm laser (CI_95_ = 21.14, 30.81), confirming the suitability of the SiMPull surface for studying synaptosomes, without significant background (noise) signal. The next question concerned the fluorescence properties of the synaptosomes. Since most brain tissue samples are auto-fluorescent^26^, which may confound the signal acquired from immunolabeling, we aimed to test if the synaptosomes also fluoresce under light excitation, without any fluorophore labelling. No significant signal was detected from the mouse (CI_95_ = 21.14, 30.81) or human (CI_95_ = 2.83, 9.18) synaptosomes in the absence of CellMask. Binding properties of CellMask to the SiMPull surface was also a concern, since unspecific binding could confound the signal. To test this, we compared the signal acquired with or without synaptosomes, in the presence of CellMask and observed significantly less signal from CellMask in the absence of synaptosomes (CI_95_ = 25.64, 36.95). Collectively, these results showed that neither the SiMPull surface, nor the synaptosomes emit any signal on their own and a label (CellMask) is necessary to identify the synaptosomes. Moreover, CellMask does not show unspecific-binding to the SiMPull surface. As such, the SynPull methodology with CellMask is well suited to study synaptic fragments.

**Figure 2.**
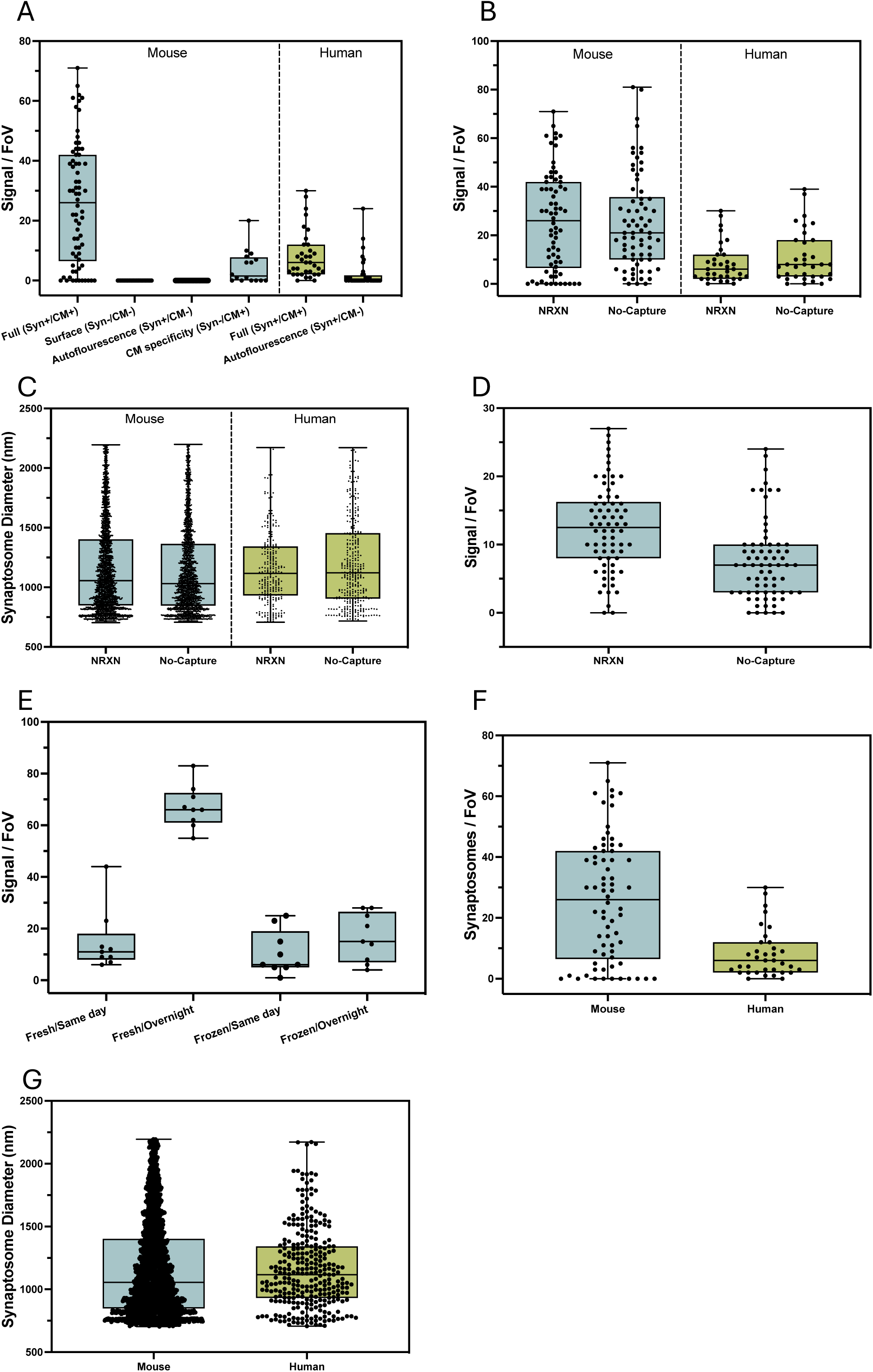
Control studies. (**A**) Neither the SiMPull surface nor the synaptosomes are auto-fluorescent, and CellMask does not non-specifically bind to the surface. (**B**) The number and (**C**) diameter of the CellMask-positive signal does not differ between the anti-neurexin captured and un-captured samples, however (**D**) Bassoon signal is higher in the anti-neurexin captured samples, indicating higher synaptosome localisation on the SiMPull surface when the anti-neurexin capture is used. (**E**) Freezing the synaptosomes decreases the signal, while overnight incubation on the SiMPull surface at 4℃ increases the signal. (**F**) While the same mass of mouse brain tissue yield a higher number of synaptosomes (**G**) the diameter of the synaptosomes does not differ between human and mouse brain tissue.

To study the efficiency of capturing the synaptic fragments with an anti-NRXN1 antibody, we quantified the CellMask signal and measured synaptosome size, as well as the bassoon signal, with and without a capture antibody. There was no significant difference between CellMask signal for mouse (CI_95_ = -6.32, 7.06) or human (CI_95_ = -7.06, 1.86) samples (**Figure 2B**), and the average synaptosome diameter did not differ considerably for mice (CI_95_ = 0.06, 45.33) or humans (CI_95_ = -96.94, 8.32; **Figure 2C**). However, the bassoon co-localisation with CellMask was significantly higher in the samples captured with the anti-NRXN antibody (CI_95_ = 2.90, 6.98; **Figure 2D)**, suggesting the specific capturing of synapses and not liposomes when a capture antibody is used.

We concluded the control experiments by determining the effects of varying incubation and storage conditions on the synaptosomes and comparing the tissue samples from mouse models of neurodegeneration and human post-mortem pre-frontal cortex tissue samples. While freezing the sample after preparing the synaptosomes decreased the yield significantly (CI_95_ = 41.74, 59. 37), incubating the sample on the surface overnight at 4℃ increased the signal (CI_95_ = 41.95, 62.50), suggesting that the optimal condition for capturing and imaging synaptosomes is to use them fresh with an overnight incubation (**Figure 2E**). Even though mouse and human synaptosome samples were prepared from a similar amount of tissue (∼300 mg), a significantly higher number of synaptosomes were obtained from the mice (CI_95_ = 12.21, 23.22; **Figure 2F**), but synaptosome size did not differ between the species (CI_95_ = -39.96, 39.75; **Figure 2G**).

### Aggregates in human brain samples

After establishing the best SynPull assay conditions we compared the length, area, and eccentricity of the ɑSyn, Aβ and AT8-positive tau aggregates from post-mortem PD, AD, and control orbitofrontal cortex samples using *d*STORM (**see Table 1 for statistical tests**). Synaptosomes within the field of view were identified by the CellMask signal and aggregates within the borders of the synaptosomes were defined as synaptic aggregates, meanwhile the ones still captured on the SiMPull surface yet located outside a synaptosome were defined as extra-synaptic aggregates. Overall, AT8-positive tau formed the longest aggregates, with an average length close to 150 nm, which were ∼50 nm longer than the aggregates formed by ɑSyn and Aβ. Notably, the Aβ and tau aggregates were always characterised in the same samples.

**Table 1.**

|  |  |  |  |  |
| --- | --- | --- | --- | --- |
| Human post-mortem pre-frontal cortex | Alpha-synuclein<br>(PD brain) | Synaptic aggregate count | $t_{3.12} = 1.31, p = 0.28; CI_{95} = -0.78, 0.32$ | Fig. 3A |
| | | Extra-synaptic aggregate count | $t_{5.08} = 0.89, p = 0.41; CI_{95} = -665.31, 322.31$ | Fig. 3A |
| | | Aggregate length | $AIC = 18837, F = 20.81, p < 0.001, CI_{95} = 44.51, 111.62$ | Fig. 3B |
| | | Aggregate area | $AIC = 36931, F = 16.64, p < 0.001, CI_{95} = 964.13, 2748.74$ | Fig. 3C |
| | | Aggregate eccentricity | $AIC = 8658.8, F = 62.33, p < 0.001, CI_{95} = 0.05, 0.08$ | Fig. 3D |
| | | Synaptosome radius | $t_{187.12} = 5.53, p < 0.001; CI_{95} = 126.94, 267.56$ | Fig. 3E |
| | Beta-amyloid<br>(AD brain) | Synaptic aggregate count | $t_{5.32} = 0.21, p = 0.845; CI_{95} = -0.09, 0.07$ | Fig. 3F |
| | | Extra-synaptic aggregate count | $t_{3.10} = 1.61, p = 0.202; CI_{95} = -83.72, 262.22$ | Fig. 3F |
| | | Aggregate length | $AIC = 4077.8, F = 6.96, p = 0.008, CI_{95} = 2.97, 20.31$ | Fig. 3G |
| | | Aggregate area | $AIC = 4109.5, F = 39.81, p < 0.001, CI_{95} = 88.98, 169.43$<br>$AIC = 8537.7, F = 16.07, p < 0.001, CI_{95} = 3717.87, 10860.18$ | Fig. 3H |
| | | Aggregate eccentricity | all $p > 0.05$ | Fig. 3I |
| | | Synaptosome radius | $t_{134.9} = 2.45, p = 0.016; CI_{95} = 17.81, 166.99$ | Fig. 3J |
| | AT8+ Tau<br>(AD brain) | Synaptic aggregate count | $t_{3.17} = 5.63, p = 0.010; CI_{95} = 1.65, 5.68$ | Fig. 3K |
| | | Extra-synaptic aggregate count | $t_{3.01} = 13.98, p = 0.001; CI_{95} = 18304.10, 29079.40$ | Fig. 3K |
| | | Aggregate length | $CI_{95} = 43.67, 55.23$<br>$CI_{95} = -100.05, 83.28$ | Fig. 3L |
| | | Aggregate area | all $p > 0.05$ | Fig. 3M |
| | | Aggregate eccentricity | $CI_{95} = 0.04, 0.07$<br>$CI_{95} = -0.11, 0.09$ | Fig. 3N |
| | | Synaptosome radius | $p > 0.05$ | Fig. 3J |

The majority of the ɑSyn, Aβ, and AT8-positive tau aggregates were extra-synaptic, with relatively few synaptosomes containing aggregates. For ɑSyn and Aβ, the number of aggregates inside or outside the synaptosomes did not differ between the PD/AD and control brains (**Figure 3A/F**), however the AD brain contained significantly more AT8-positive tau aggregates, both inside (factor of ∼4.5) and outside the synaptosomes (factor of ∼3,000; **Figure 3K**). Meanwhile, the length (160nm vs 80nm; **Figure 3B**) and area (2640nm^2^ vs 738nm^2^; **Figure 3C**) of the synaptic ɑSyn aggregates from the PD brain were bigger than the extra-synaptic aggregates from the PD and control brain. For Aβ, once again the synaptic aggregates were longer than the extra-synaptic ones (141nm vs 83nm). However, the aggregates from the control brain had a size distribution with a longer mean (**Figure 3G**), although there were very few synaptic aggregates in the control brain, they were the longest and largest of the four conditions with an average length of 295nm and average area of 7600nm^2^ (**Figure 3H**). On the other hand, the AT8-positive tau aggregates from the same samples did not show a length difference based on AD status inside the synapse, but the extra-synaptic aggregates were longer in the AD brain, compared to the controls (150nm vs 100nm; **Figure 3L**), yet the area of the AT8-positive tau aggregates did not differ between the AD and control samples (**Figure 3M**).

**Figure 3.**
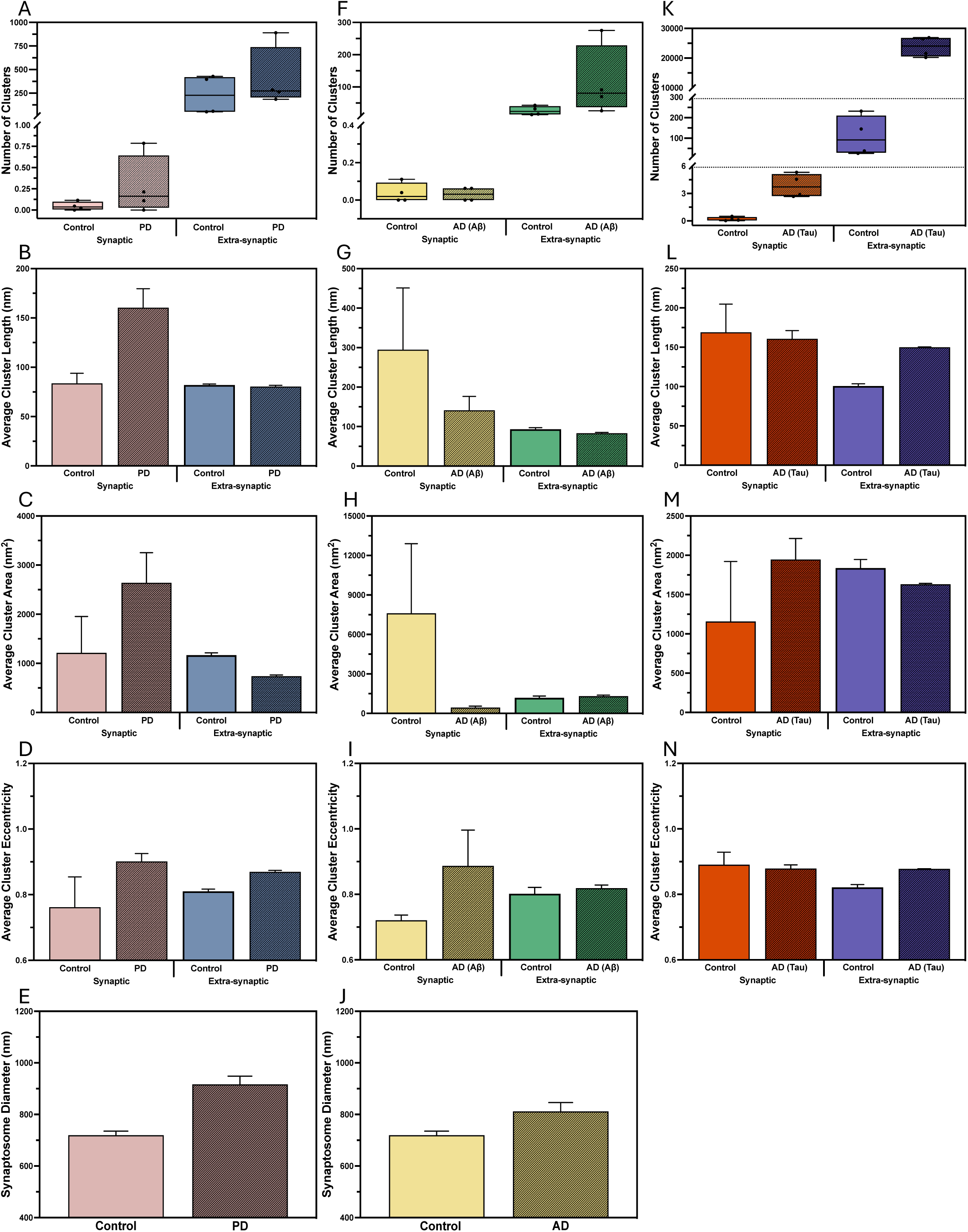
Morphological characteristics of pathological aggregates in human post-mortem brain samples. (**A, F, K**) Number of synaptic and extra-synaptic aggregates, (**B, G, L**) average aggregate length in nanometres, (**C, H, M**) average aggregate area in nanometre square, (**D, I, N**) average aggregate eccentricity (1 is flat and 0 is circular) for alpha-synuclein, beta-amyloid, and AT8-positive tau aggregates respectively from Parkinson’s and Alzheimer’s disease, and age-matched control orbitofrontal cortex samples. (**E, J**) Average synaptosome diameter. The analyses were conducted by using data from 124, 94, and 300 synaptosomes harvested from human post-mortem orbitofrontal cortex samples from Parkinson’s disease, Alzheimer’s disease brains, and control samples, respectively and analysed by linear mixed effects models and 95% confidence intervals.

The shape of the aggregates (circular or elongated) also differed between the aggregates. While the ɑSyn aggregates from the PD brain samples were more elongated (fibril-like) than the controls (**Figure 3D**), Aβ aggregates did not show a difference (**Figure 3I**), and AT8-positive tau aggregates were more elongated outside the synapse in the AD-brain samples, without a difference within the synaptic aggregates (**Figure 3N**). Lastly, the synaptosomes from the PD (**Figure 3E**) and AD (**Figure 3J**) brain samples on average had greater radius than those from control samples.

Collectively, these results are suggesting that more fibrillar aggregates are present in the PD brain samples, with larger aggregates accumulating in the enlarged synapses in PD, meanwhile the enlarged synapses in AD brains contain smaller Aβ aggregates but a greater number of AT8-positive tau aggregates.

### Aggregates in mouse models

Following the post-mortem human brain samples, we repeated the same measures in mouse models of PD (MI2^27^), AD (APP^NL-G-F28^), tauopathy (P301S^29^), and wildtype control (C57Bl6/J) at 6-months of age (**see Table 2 for statistical tests**). The aggregates formed in the mice were smaller than the human brain samples, with an average length under 100 nm for all species, and there was an overall trend of longer aggregates in the synapse.

**Table 2.**
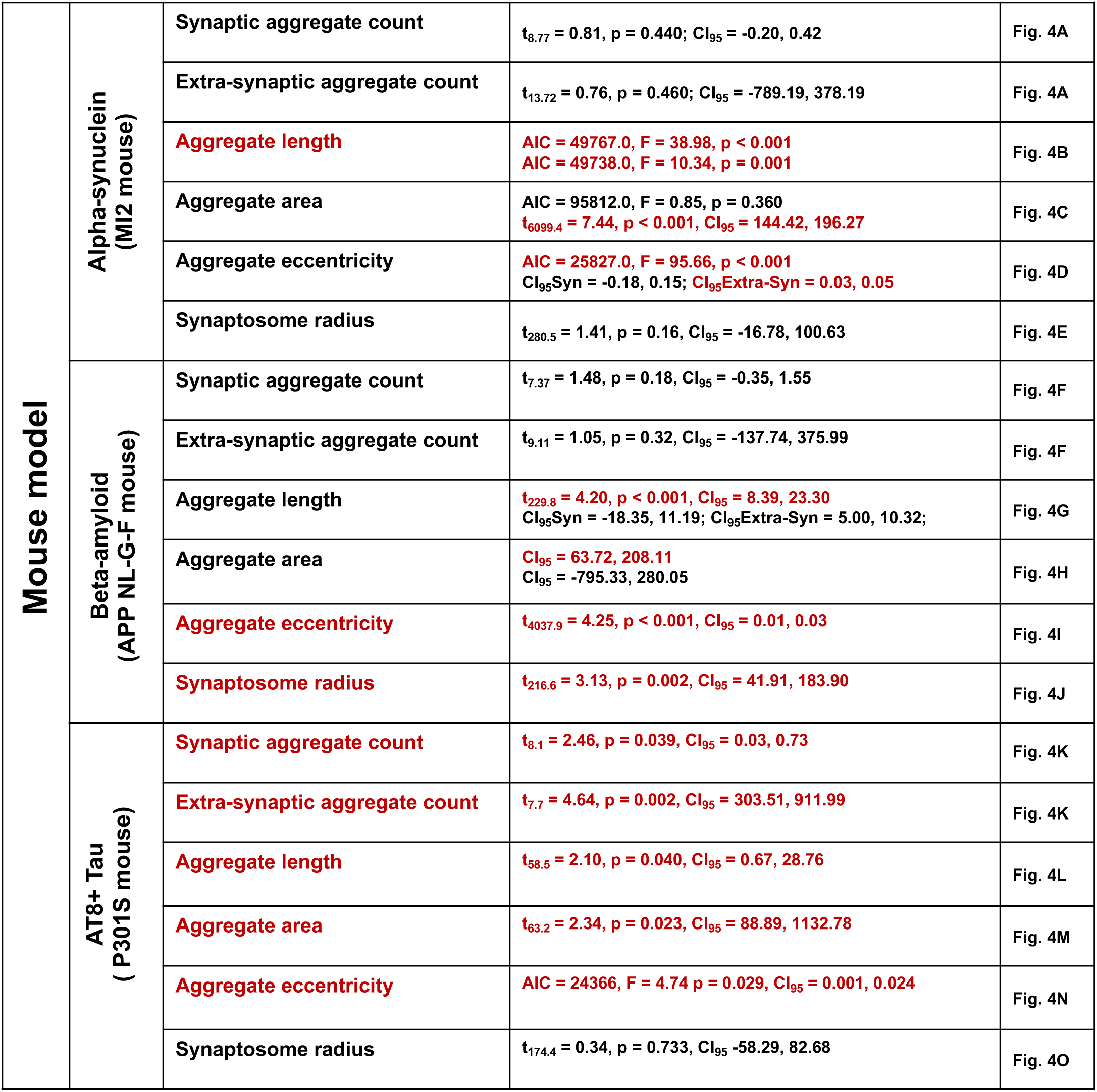

There were prominent parallels between the mouse models and the human brain samples. For instance, the majority of the ɑSyn, Aβ, and AT8-positive tau aggregates were found in the extra-synaptic fractions in the mouse samples as well. Moreover, the number of ɑSyn and Aβ aggregates did not differ between the transgenic (MI2 and APP^NL-G-F^) and wildtype control mouse brains (**Figure 4A/F**), but the P301S mice had significantly more AT8-positive tau aggregates than the controls both in the synaptosomes and the extra-synaptic samples (**Figure 4K**). While the ɑSyn aggregates from the MI2 mice were longer than the ones in the control mice (∼85nm vs 100nm), the synaptic ɑSyn aggregates were longer than the extra-synaptic ones regardless of genotype (∼110nm vs 80nm; **Figure 4B**). Similarly, the APP^NL-G-F^ mice had longer Aβ aggregates than the controls (∼83nm vs 75nm), but the difference reached statistical significance only for the extra-synaptic aggregates. Meanwhile, the synaptic Aβ aggregates were longer than the extra-synaptic ones, regardless of genotype (∼96nm vs 80nm; **Figure 4G**). A similar trend was observed for the AT8-positive tau aggregates as well, as the synaptic aggregates were significantly longer than the extra-synaptic ones (∼96nm vs 76nm), regardless of genotype (**Figure 4L**).

**Figure 4.**
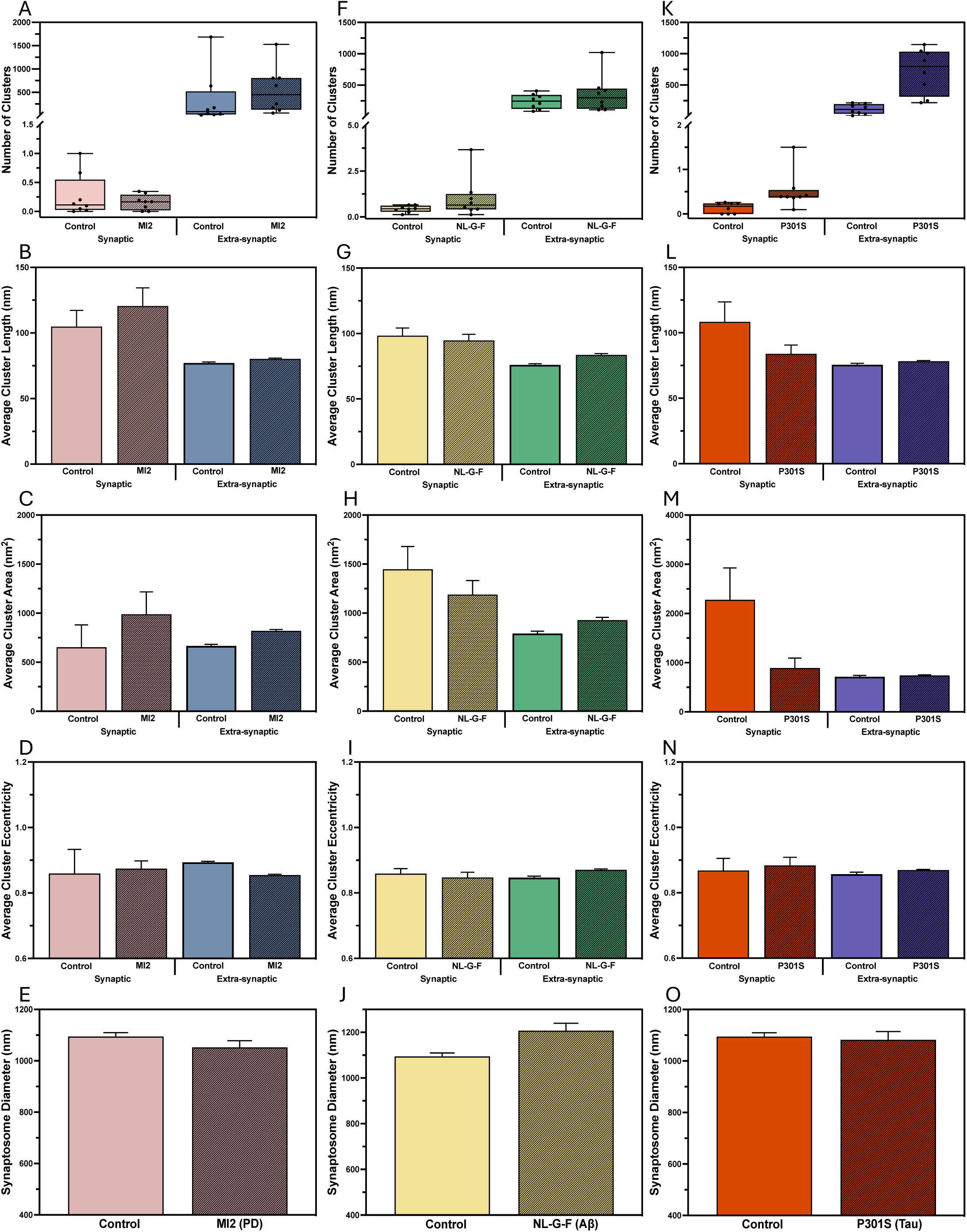
Morphological characteristics of pathological aggregates in MI2, APP^NL-G-F^, P301S, and C57Bl/6J mouse brain samples. (**A, F, K**) Number of synaptic and extra-synaptic aggregates, (**B, G, L**) average aggregate length in nanometres, (**C, H, M**) average aggregate area in nanometre square, (**D, I, N**) average aggregate eccentricity (1 is flat and 0 is circular) for alpha-synuclein, beta-amyloid, and AT8-positive tau aggregates respectively from MI2, APP^NL-G-F^, P301S, and C57Bl/6J mouse brains. (**E, J, O**) Average synaptosome diameter. The analyses were conducted by using data from 160, 149, 119, and 520 synaptosomes harvested from MI2, APP^NL-G-F^, P301S, and C57Bl/6J mouse brains, respectively and analysed by linear mixed effects models and 95% confidence intervals.

Similar to ɑSyn aggregate length, the average aggregate area was also larger for the MI2 mice, compared to the controls, yet there was no area difference between the synaptic and extra-synaptic aggregates (**Figure 4C**). While the area of the synaptic Aβ aggregates did not show a genotype difference, the APP^NL-G-F^ mice had larger aggregates in the extra-synaptic fragments (**Figure 4H**). On the other hand, the synaptic AT8-positive tau aggregates were significantly larger than the extra-synaptic ones, regardless of genotype (**Figure 4M**).

The trends for aggregate shape differed considerably between the species; while the extra-synaptic ɑSyn aggregates from the MI2 mice were more circular than the ones from the control mouse brain (**Figure 4D**), the Aβ and AT8-positive tau aggregates were more elongated in the brains of the APP^NL-G-F^ (**Figure 4I**) and P301S mice (**Figure 4N**) respectively. Lastly, there was no genotype difference between the synaptosomes size from MI2 and P301S mouse brains compared to the controls (**Figure 4E/O**), but the synaptosomes from the APP^NL-G-F^ mice had a greater radius compared to the ones from the control mice (**Figure 4J**).

The abundance of AT8-positive tau aggregates in human and mouse synaptosomes allowed us to perform additional statistical analyses, specifically to investigate if the appearance of each individual aggregate is a random event or if the presence of one aggregate increases the chance of finding another in the same synaptosome. If indeed the appearance of aggregates were random, the distribution of aggregates per synaptosome should follow a Poisson distribution. Based on the fraction of synaptosomes that contain no aggregates, we should then be able to predict the fractions of synaptosomes that have one aggregate, two aggregates, and so on. In **Figure 6**, we show the experimentally measured distribution as well as this prediction (solid grey line). We also show the Poisson distribution based on the average number of aggregates per synaptosomes. If the data are Poisson distributed, those two predictions would match. Indeed, we find that this is the case for both mouse datasets (**Figures 6C&D**), as well as the control human dataset (**Figure 6A**). However, the distribution of aggregates per synaptosome obtained from the human AD samples clearly deviates from a Poisson distribution (**Figure 6B**). The two predictions, based on the average number or the fraction of synaptosomes without aggregates, do not match with each other or with the experimental data and there is a significantly higher fraction of synaptosomes with multiple aggregates than would be expected for a random Poisson distributed event. In other words, if a synaptosome has one AT8-positive tau aggregate, it is more likely to have another -than would be expected if the appearance of aggregates were independent events.

**Figure 5.**
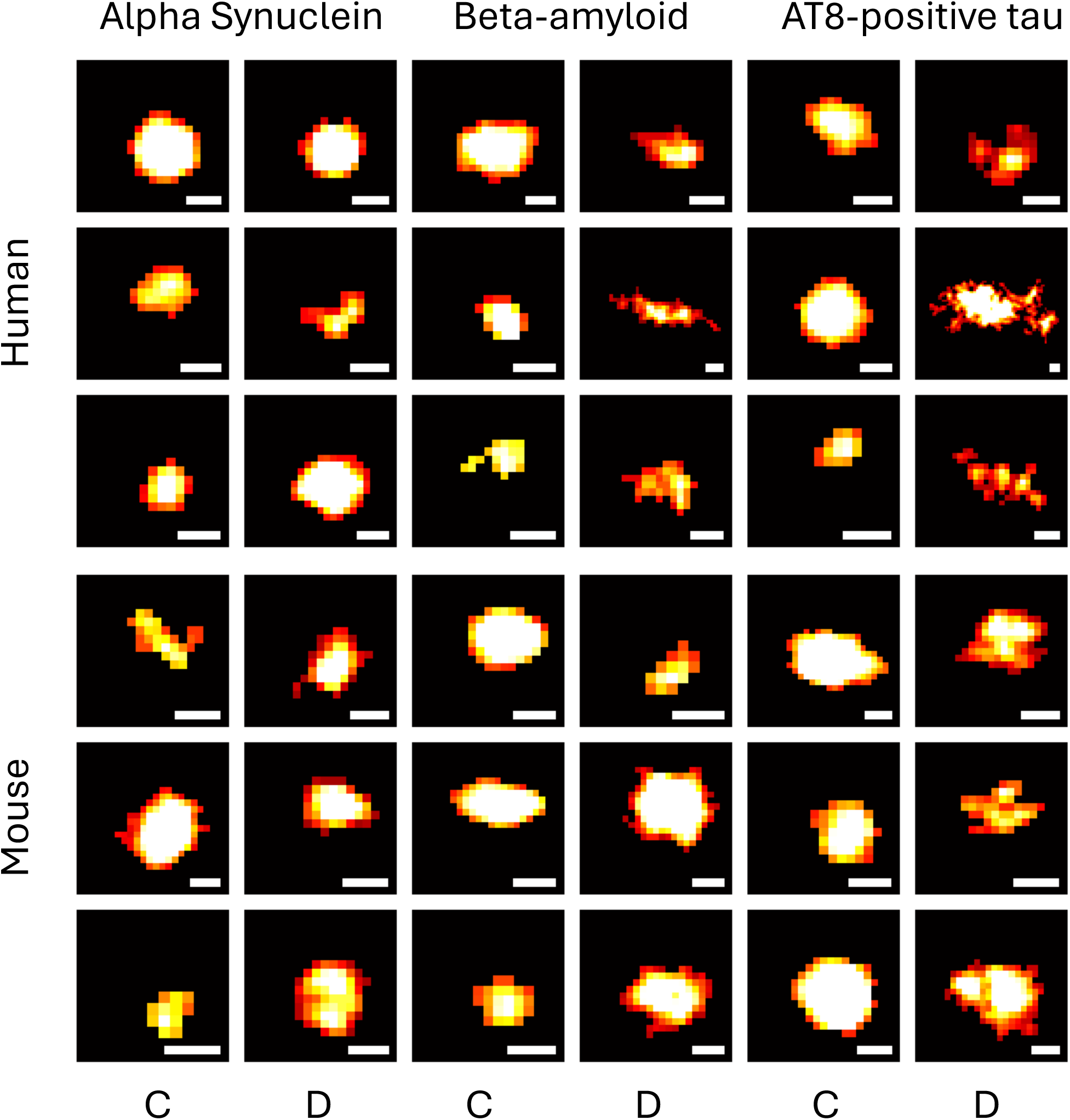
Representative images of alpha-synuclein, beta-amyloid, and AT8-positive tau aggregates from human and mouse brain samples acquired by from *d*STORM microscopy and reconstructed by ACT software. Scale bars are 50 nm.

**Figure 6.**
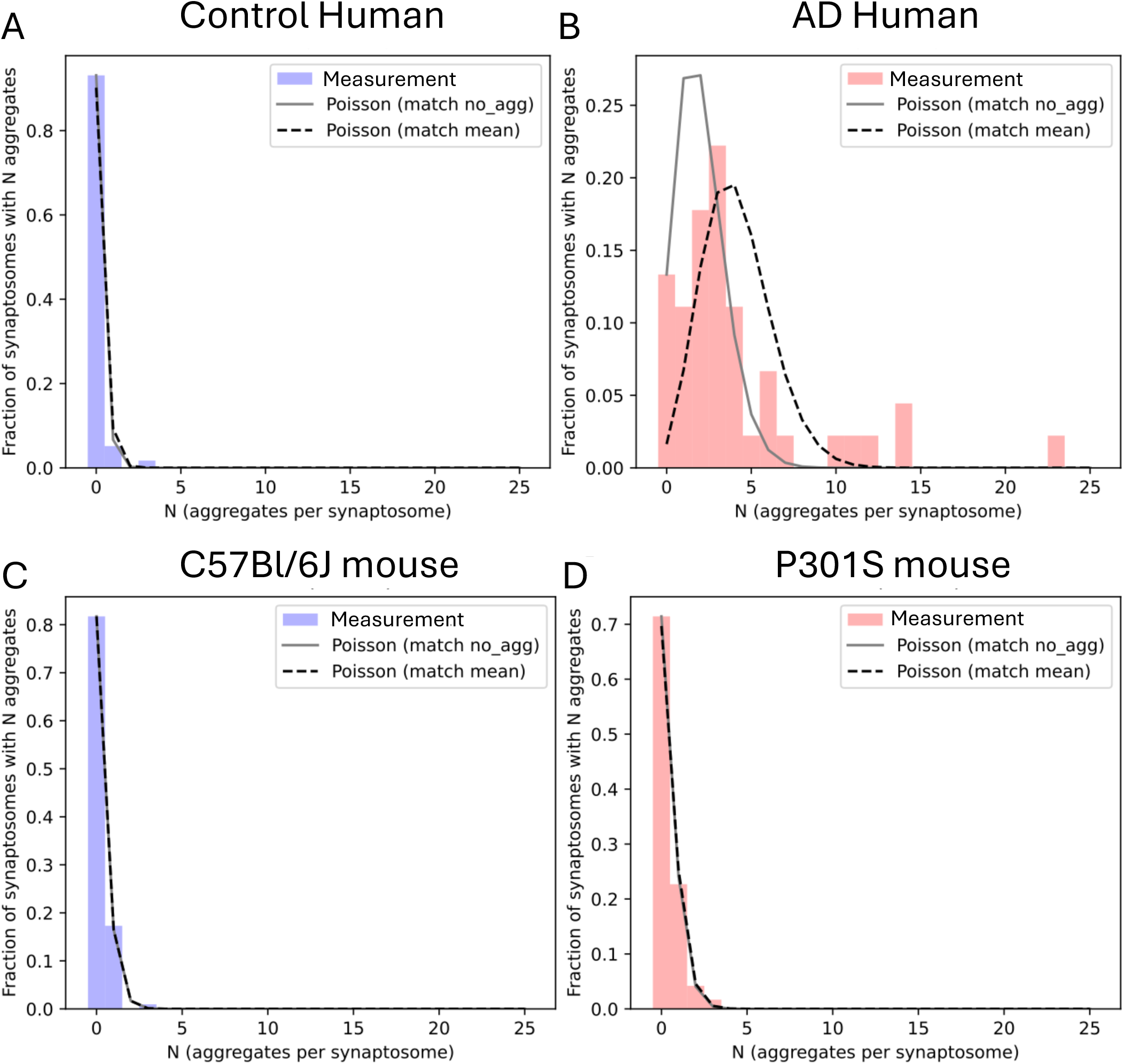
Aggregate per synaptosome distributions from experiment and theory. Experimentally measured aggregate per synaptosome distributions (bars), for (**A,B**) human and (**C, D**) mouse samples. The predicted Poisson distribution, based on either matching the fraction of synaptosomes without aggregates (solid grey line) or the average number of aggregates per synaptosome (dashed black line) is shown overlaid. For all except the data from human AD samples (**B**) the 2 predicted distributions overlap with each other and match the data. For the human AD samples, there are more synaptosomes with higher numbers of aggregates than would be expected based on the total fraction of synaptosomes that contain aggregates (compare bars to solid grey line), indicating that the appearance of individual aggregates are not independent events as in the other 3 datasets.

## 3. Discussion

Synaptic dysfunction is one of the core pathologies seen in AD and PD, promoting cognitive and behavioural deficits. Interestingly, Aβ, tau, and ɑSyn all have physiological roles in the synapse but their pathological aggregation and accumulation causes synaptic dysfunction and loss^30,31^. Here, we developed SynPull as a novel method for characterising soluble aggregates inside the synapses by combining single-molecule pulldown, *d*STORM super-resolution imaging, and advanced computational analysis tools and applied them to post-mortem brain samples from AD and PD patients, as well as mouse models.

We used the pre-synaptic protein neurexin 1 to reliably capture the synaptosomes. Since the fractured membrane particles can also fuse and form liposomes, using a synaptic protein to capture the bassoon-positive synaptosomes enabled us to reliably target the liposomes containing the synaptic compartment (**Figure 2D**). Then, we used image analysis tools to identify the synaptosomes based on their size, determined by the CellMask signal (**Figure 2G**). Lastly, we used *d*STORM to super-resolve the aggregates inside and outside the synaptosomes, allowing the morphological characterisation of individual soluble aggregates with a resolution-limit of 20 nm (**Figure 5**). Importantly, these methods are not limited to synaptosomes from human brains and mouse models of neurodegeneration; a broad range of pathologies and/or developmental stages can be reliably studied with high specificity, by using appropriate capture antibodies.

Even though the primary aim of this work was to develop a novel method to study small-soluble aggregates in the synapse, by studying post-mortem human disease brain samples and mouse models, we could make biological observations. Regardless of aggregate type, most of the aggregates were extra-synaptic with relatively few synaptosomes containing one or more aggregates (**Figure 7**). While the least common aggregate type in the synapse was Aβ, AT8-positive tau was the most common aggregate. This observation is interesting since both Aβ and tau aggregates were studied in the same human sample, obtained from the same AD brain, and it is in-line with the findings of Colom-Cadena *et al*., who found abundant oligomeric tau accumulation in the AD brain, even in regions without robust tau tangle pathology^16^.

**Figure 7.**
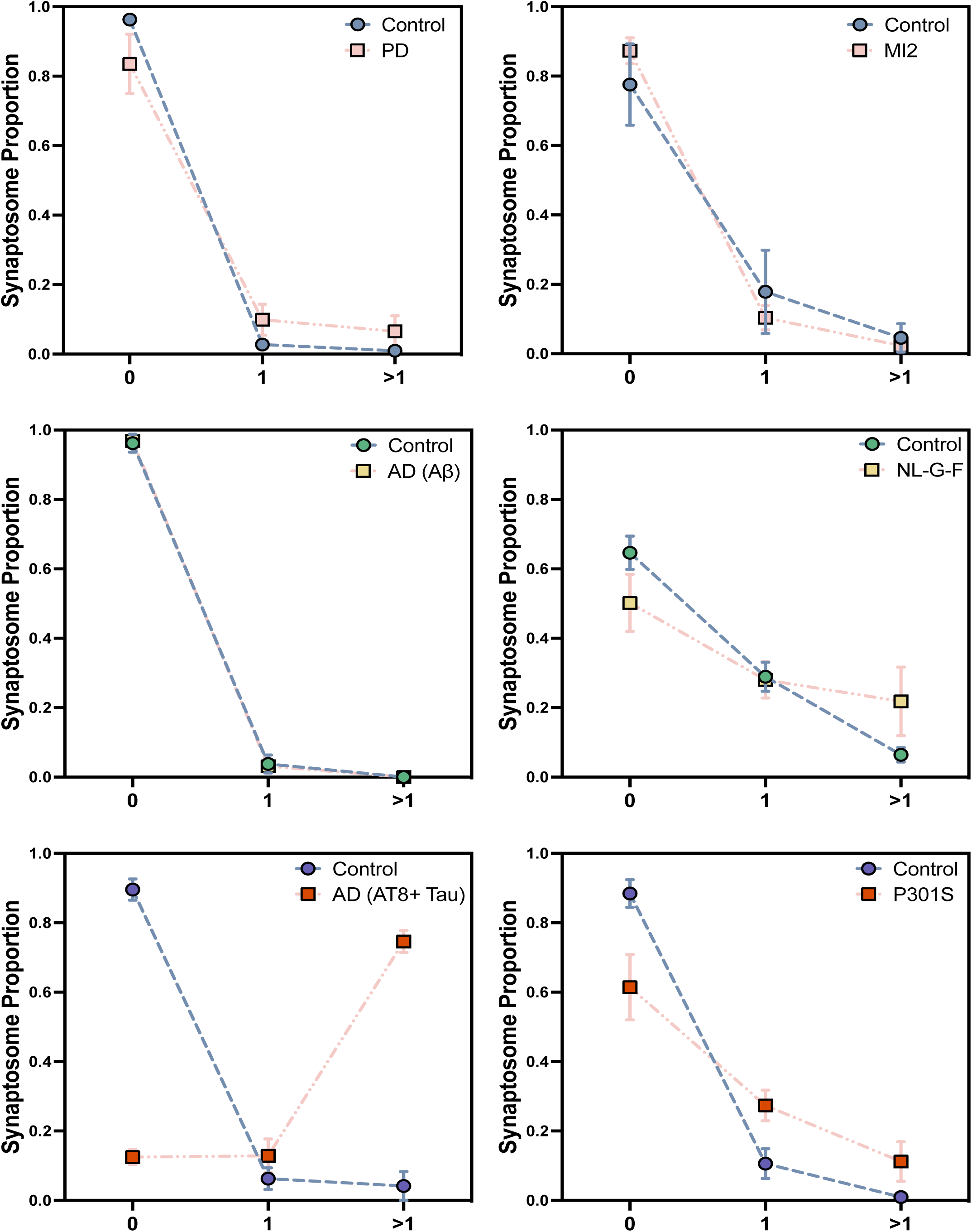
The proportion of synaptosomes with zero, one, and greater than one alpha-synuclein (top row), beta-amyloid (middle row), and AT8-positive tau (bottom row) aggregates from post-mortem human (left column) and mouse (right column) brain samples.

The Aβ and ɑSyn aggregates inside the synaptosomes were larger than the extra-synaptic ones, indicating the presence of a specific population of aggregates accumulating in the synapses. Moreover, the synaptosomes in human brains from the AD and PD patients were larger than those from controls. This has been previously observed in AD brain samples and suggested to reflect a compensatory mechanism against synaptic loss^32^. This observation of potential enlarging of the synapse to make up for synaptic loss also shows a possible limitation of post-mortem synaptosome studies: since synaptic loss is common in both AD and PD, the synaptosomes available for harvesting in the post-mortem tissue may reflect a survival bias.

The presence of Aβ in the synapse and its physiological role has been long studied. While monomeric Aβ at physiological levels seems to be required for synaptic function^33^, oligomeric aggregates disrupt both synapse formation^34^ and function^35^. Interestingly, monomeric Aβ has been shown to interact with the soluble oligomeric aggregates and reduce their toxicity^36^. On the other hand, the presence of Aβ has been shown to translocate tau to the synapse^13^. Our results agree with this, since the synaptic AT8-positive tau aggregates were significantly higher in the AD brain. Since the synaptosomes were prepared from the same brain samples for Aβ and tau analyses, our results are also suggesting that tau is significantly more common in the orbito-frontal cortex synapses at later AD stages. Our data modelling also revealed that that presence of AT8-positive tau aggregates in the synapse is not a random event and having one aggregate in the synapse increases the chance of having multiple aggregates (**Figures 6&7**). Mechanistically, there are two possible interpretations of this observation: (1) the appearance of the first aggregate makes the appearance of the second more likely, possibly by self-replication of the first aggregate. (2) A confounding factor means that one subset of synapses contains a much higher average number of aggregates than the other subset. This could be, for example, the subset of synapses that is located close to a Aβ plaque, or the subset of synapses that is connected to cells which contain a tau tangle. The fact that this behaviour is only observed in the data from human AD samples, but not in those from the P301S mice suggests that Aβ may be an important factor in synaptic AT8-positive tau aggregation, even though Aβ itself does not aggregate in the synapse. Alternatively, it may also be due to the differences in the rate of pathology in humans with AD, which takes decades and the mouse model, in which the pathological tau expression begins early in life and is promoter-regulated.

Interestingly, unlike Aβ and ɑSyn, the synaptic tau aggregates did not differ in size from the extra-synaptic ones. However, it must be noted that all aggregates were harvested by the synaptoneurosome preparation protocol, which only yields a sub-population of the soluble aggregates in the tissue samples. Indeed, the length distribution of tau aggregates in the homogenised AD brain samples were shown to be slightly smaller compared to our previous work, which studied all the soluble AT8-positive tau aggregates harvested through homogenisation of the brain tissue^37^. On the other hand, the greatest size difference based on disease condition and synaptic localisation was observed in the ɑSyn aggregates, studied in PD brain samples. Similar to Aβ the (patho)physiological role of ɑSyn in the synapse is believed to be determined by its aggregation state: in its native state, ɑSyn is membrane-bound and interacts with synaptobrevin-2 to facilitate SNARE complex formation and neurotransmitter release^38^. However, when detached from the vesicle membrane, ɑSyn can start to aggregate, which leads to loss of function as well as gain of toxic function^30^. Lacking a transmembrane domain, ɑSyn easily dissociates from membranes, which led to the lack of its observation in earlier synaptic isolations^39^. Nevertheless, more recent studies with highly-sensitive methods determined the presence of ɑSyn in synaptosomes as well as synaptic vesicles^40,41^, showing the utility of highly-sensitive methods^42^.

While these results are showing the utility of SynPull to study sub-diffraction limit pathological aggregates in synapses, which are invisible to most other imaging techniques, it has some limitations that can be addressed in future work. While synaptosomes are three-dimensional structures, the imaging within the TIRF field is two-dimensional, disabling the exact localisation of the aggregates within the synaptosome. Moreover, the relatively low yield of human brain for synaptosomes (**Figure 2F**) requires manual scanning of the surface to locate and image the synaptic fragments, which is time consuming. Further optimisation of the harvesting to enrich for synaptosomes would be beneficial to accelerate the imaging process, by allowing the usage of automated scanning. Lastly, while the results we present here show the utility of SynPull, they need to be further verified in larger sample cohorts, including multiple brain regions and disease stages, as well as different antibodies specific to different Aβ fragments, phosphorylation of tau at other regions, and ɑSyn with post-translational modifications.

In conclusion, here we demonstrated a novel method to study the morphology of the small-soluble aggregates in synapses, using a specific and selective capturing method, along with super-resolution microscopy, aided by specialised data processing techniques. Using these methods, we have studied Aβ, AT8-positive tau, and ɑSyn in human brain samples as well as mouse models. We have shown that tau is the most common aggregate type in the synapses in AD brain, meanwhile, even though they are much more rare, Aβ and ɑSyn aggregates in the synapse are larger than the extra-synaptic ones.

## 4. Methods

### Human and mouse brain samples

3 PD, 3 AD, and 3 age- and sex-matched (two female, one male) post-mortem human orbitofrontal cortex (Brodmann areas 10-11) samples were used alongside brain samples from MI2, APP^NL-G-F^, P301S, and C57Bl/6J mice. The human disease brain samples were selected from Lewy body Braak stages 4/5 for PD^43^ and neurofibrillary tangle Braak stages 5/6 for AD^44^. Due to the extensive dopaminergic input to this region from the midbrain, the orbitofrontal cortex is involved in the decision making and reward processing deficits seen in PD^45,46^. Similarly, extensive neurofibrillary tangle pathology is observed in the orbitofrontal cortex of AD patients, especially in Layers 3 and 5^47^. Brain samples were snap frozen and stored frozen at –80℃ until synaptosome preparation.

The MI2 mice express truncated 1-120 αSyn under the control of the tyrosine hydroxylase promoter in the C57Bl/Ola background without any endogenous αSyn expression, resulting in the selective expression of aggregation-prone C-terminal truncated αSyn in dopaminergic neurons, associated with synaptic aggregation and dysfunction^27^. Three homozygous, 6-month-old, male mice were used in this study, an age which dopaminergic neuronal dysfunction is observed. Due to the anatomically limited expression of the transgene to the dopaminergic neurons, brains were sliced into two parts, separating the midbrain and the dorsal striatum from the rest of the cerebrum.

The APP^NL-G-F^ mice express human amyloid precursor protein (APP) with the Arctic, Swedish, and Iberian mutations under the murine APP promoter and show plaque pathology starting at 2 months of age, which increases up to 7 months of age^28^. Synaptic loss along with cognitive impairment has been observed in this model by 6-months of age^28^. Three homozygous, 6-month-old, male mice were used in this study.

The P301S mice express human 4R/0N tau with the P301S mutation under the neuronal portion of the murine Thy1 promoter, resulting in tau tangle pathology, neuronal loss, and behavioural deficits by 4-months of age^29,48^. Three male mice at 6-months of age were used in this study.

### Synaptosome preparation

The synapto(neuro)some preparation method developed by Tara Spires-Jones and colleagues^49^ was used in these experiments. In brief, a full mouse cerebrum (or human tissue of similar size of 300-400 mg) was initially homogenised with 1 mL, ice-cold homogenisation buffer, containing 25 mM HEPES (pH 7.5), 120 mM NaCl, 5 mM KCl, 1 mM MgCl2, and 2 mM CaCl2, dissolved in HPLC-grade water, using a 2 mL Dounce homogeniser (Cambridge Scientific, Cat. 40401). Then the homogenate was first filtered through an 80 µm nylon filter (Millipore, Cat. NY8002500; 25 mm filter holder: PALL, Cat. 4320) to remove tissue debris, followed by a second filtration with a 5 µm filter (Millipore, Cat. SMWP04700), to remove the organelles and the nuclei. Then the homogenate was collected in a 1.5 ml Lo-bind Eppendorf and centrifuged at 1000 g for 5 minutes at 4℃. Because this preparation yields synaptic fragments containing the complete pre-synapse as well as parts of the post-synaptic compartment, the synaptic fragments prepared using this method are termed synaptoneurosomes^24,49^. A total of 1466 synaptosomes (124 from PD, 94 from AD, 300 from control human brain samples along with 160 from MI2, 149 from APP^NL-G-F^, 119 from P301S, and 520 from C57Bl/6J control mice) were harvested and studied.

### Transmission electron microscopy (TEM)

Samples were fixed with a mixture containing 2% glutaraldehyde and 2% formaldehyde in 0.05 M sodium cacodylate buffer (pH 7.4) containing 2 mM CaCl_2_ overnight at 4℃. After the fixation, samples were washed five times with 0.05 M sodium cacodylate buffer (pH 7.4) and osmicated in 1% osmium tetroxide and 1.5% potassium ferricyanide in 0.05 M sodium cacodylate buffer (pH 7.4) and osmicated in 1% osmium tetroxide and 1.5% potassium ferricyanide in 0.1 M sodium cacodylate buffer (pH 7.4) for 3 days at 4℃. All samples were then washed five times in deionised water (DIW) and treated with 0.1 % (w/v) thiocarbohydrazide in DIW for 20 minutes at room temperature in the dark. After washing five times with DIW, samples were osmicated a second time for 1 hour at room temperature (2% osmium tetroxide in DIW). After washing five times with DIW, samples were block stained with uranyl acetate (2% uranyl acetate in 0.05 M maleate buffer -pH 5.5) for 3 days at 4℃. Samples were washed five times in DIW and then dehydrated in a graded series of ethanol (50%, 70%, 95%, 100%, 100% dry) and 100% dry acetonitrile, three time in each for at least 5 minutes. Samples were infiltrated with a 50/50 mixture of 100% dry acetonitrile/Quetol resin (without BDMA) overnight, followed by 3 days in 100% Quetol (without BDMA). Then, samples were infiltrated for 5 days in 100% Quetol resin with BDMA, exchanging the resin each day. The Quetol resin mixture is: 12 g Quetol 651, 15.7 g NSA, 5.7 g MNA and 0.5 g BDMA (all from TAAB). Samples were placed in embedding moulds and cured at 60℃ for 3 days.

90 nm thick TEM sections were cut by an ultramicrotome (Leica Ultracut) and placed on 300 mesh bare copper grids. Samples were imaged in a Tecnai G2 TEM (FEI/ThermoFisher) run at 200 keV accelerating voltage using a 20 μm objective aperture to improve contrast; images were acquired using an AMT digital camera.

### SiMPull coverslip preparation

The SiMPull coverslips were prepared as previously described^50^. In brief, glass coverslips were first cleaned with argon plasma and then coated with a 1:1 mixture of Rain-X liquid and isopropanol. Once the mixture was fully evaporated, the slides were stored in a dark cabinet at room temperature.

### Capturing the synaptosomes on SiMPull

SiMPull coverslips were stored in a dark and dry environment in room temperature until used. First, the wells were washed twice with phosphate buffered saline (PBS). Then, neutravidin dissolved in PBS at 0.2 mg/ml was incubated for 10 minutes, followed by two more PBS washes. Next, a Pluronic F-127^™^ polymer (Biotium, Cat. 59005) dissolved in PBS (1:10 by volume and with 0.2µm filtering) was incubated on the surface for 90 minutes, followed by two washes with PBS with 0.05% Tween (PBST). A biotinylated polyclonal anti-neurexin 1 (NRXN1) antibody (abcam, Cat. ab222806) at 10 nm concentration (in PBST) was added to the surface for 10-minutes, followed by two washes with PBST. Finally, 10 µL of synaptosome sample was added to the surface and incubated overnight at 4℃.

### Staining the synaptosomes

An orange CellMask^™^ plasma membrane stain (Thermo Fischer Scientific, Cat. C10045) was used to label the synaptosomes. Following the overnight incubation of the samples, the surface was washed twice with PBST and a CellMask solution (1:6000 by volume in PBS) was added to the surface for 10 minutes, followed by four rounds of washes with PBS.

### Immunolabeling the synaptic proteins

In order to study pre- and post-synaptic marker proteins, as well as Aβ aggregates inside the synaptosomes, they were fixed, and their membranes were permeabilised. First, they were incubated in a formaldehyde (Pierce^™^ 16% Formaldehyde (w/v), Methanol-free, Thermo Fischer Scientific, Cat. 28906) solution (4:100 by volume in PBS) for 10-minutes, followed by quenching with 20 mM glycine for another 10-minutes. Then the samples were washed twice with PBS and incubated in a permeabilising solution consisting of Triton-X (Sigma-Aldrich, Cat. X100-5ML) dissolved in PBS (1 in 100,000 by volume) for 5-minutes, followed by two washes with PBST. Then, the monoclonal anti-bassoon (abcam, Cat. ab82958) antibody was used to identify the pre-synaptic compartments, while the post-synapse was identified using anti-postsynaptic density (PSD)95 (abcam, Cat. ab18258). The antibodies were labelled with Alexa Fluor^™^ 488 and 647 NHS ester (Thermo Fischer Scientific, Cat. A20000 and A37573, respectively). The antibodies were incubated on the surface at 5 nM concentration for 20 minutes.

Bassoon was used in these experiments as a pre-synaptic marker due to its selective immunoreactivity in the pre-synaptic boutons throughout the brain^51^. PSD95 instead was used as a marker for the post-synapse, due to its specific immunoreactivity in the post synapse.

In order to image the ɑSyn, Aβ, and AT8-positive tau aggregates, monoclonal 4B12 (Thermo Fisher Scientific; Cat. MA1-90346), 6E10 (BioLegend; Cat. 9340-02), and AT8 (Invitrogen; Cat. MN1020B) antibodies labelled with Alexa Fluor^™^ 647 NHS ester were used at 5 nM, 1 nM, and 2 nM concentration and 30 minutes, 45 minutes, and 15 minutes incubation time, respectively.

### *d*STORM imaging

Images were acquired on a custom-built single molecule localisation microscope (NanoPro) whose architecture and operation have been previously described in detail^52^. Briefly, a laser diode (PD-01229, LaserTack GmbH, Germany) emitting at 638 nm and operating at 420 mW was focused onto a 70 mm x 70 mm squared-core multimode fibre (05806-1 Rev. A, CeramOptec GmbH, Germany) agitated using a brushless DC motor (304-111, Precision Microdrives, UK) operated at <14,000 rpm. Light exiting the fibre was collimated and then focused at the back focal plane of an apochromatic, 100x, 1.49 numerical aperture objective (MRD01991, Nikon, UK). The excitation was translated to illuminate the sample under Total Internal Reflection Fluorescence (TIRF). Emitted fluorescence was then collected onto a scientific Complementary Metal Oxide Semiconductor (sCMOS) camera (O1_PRIME_BSI_EXP, Teledyne Photometrics, UK). For each field of view, 2,000 frames were acquired at an exposure time of 50 ms.

### Image processing and analysis

The acquired images were processed using the Aggregate Characterisation Tool (ACT)^53^. Briefly, images were thresholded, dilated, and eroded to segment single molecules. The location of segmented molecules was calculated using ThunderSTORM^54^, and super-resolved images were calculated by super-imposing Gaussian-blurred localisation with localisation precisions of less than 20 nm.

The CellMask images were used to distinguish the synaptosomes using a custom segmentation protocol. Before imaging the synaptosomes, images of TetraSpeck fluorescent spheres (ThermoFisher, Cat. T7279, 1:1000 in PBS) in the same field-of-view (FoV) were obtained in both the CellMask (561 nm) and antibody (640 nm) channels. These images were used to generate an affine matrix, which was then applied to the CellMask images of the synaptosomes to correct the aberration-induced misalignment between the two channels. The corrected CellMask images were firstly thresholded, and the objects of interest were identified by selecting the regions where the intensity was higher than the average plus twice the standard deviation intensity within the FOV. Small objects induced by noise were removed by erosion and dilation operations. The remaining objects were size-filtered to isolate individual synaptosomes. A similar strategy was previously employed to segregate different synaptic compartments^55–57^.

Next, a density-based clustering algorithm (DSBCAN)^58^ with two minimum points and 75 nm epsilon distance was performed on the super-resolution localisations in the 640 nm channel to cluster localisations into aggregates. Individual aggregates (as shown in **Figure 5**) were segmented using the Automated Structures Analysis Program (ASAP)^59^. Specifically, clusters with centroids within the identified synaptosomes were tagged as synaptic clusters, whereas those outside were labelled as extra-synaptic. Finally, the morphological features of these individual clusters were extracted and analysed.

### Statistical analyses

The R Project Statistical Computing version 4.2.2 (2022-10-31)-" Innocent and Trusting" was used for all statistical analyses, the graphs were generated in GraphPad Prism 7.0a for Mac OS X, and the cartoon figures were created with BioRender.com. Linear mixed effects models, ANOVA’s with Type 2 sums of squares, and 95% confidence intervals (CI) were used to determine the differences between the groups (**Tables 1 and 2**). The datasets generated and analysed during the current study are available from the corresponding author on reasonable request. The novel code generated for the analysis of the super-resolved images is provided in the supplemental material.

For data modelling and in order to produce the predictions in **Figure 7**, the Poisson distribution was used, given by the probability mass function

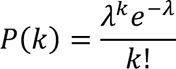

where k is the number of aggregates per synaptosome in question and λ is the average number of aggregates per synaptosome. This describes the distribution of independent events, given their average number, λ, and is expected to describe the number of aggregates per synaptosome if they appeared independently of each other, and with the same probability in each synaptosome. The solid grey line is produced by setting λ = -log(f0) where f0 is the experimentally measured fraction of synaptosomes with no aggregates. The dashed black line is produced by setting λ equal to the experimentally measured mean number of aggregates per synaptosome. The plots were produced in Python using the matplotlib and scipy packages.

## 5. Declarations

### Ethical approval

Human post-mortem brain tissue was acquired from the Cambridge Brain Bank (Cambridge University Hospitals). The Cambridge Brain Bank is supported by the NIHR Cambridge Biomedical Research Centre (NIHR203312). We gratefully acknowledge the participation of all our patient and control volunteers.

### Availability of data and materials

Data collected during these experiments and the novel code generated for image analysis will be made available upon reasonable request from the corresponding author.

### Author contributions

**S.K.** Conception and design, data collection and image analysis, manuscript preparation. **E.F.** Conception and design, data collection and statistical analysis, manuscript writing. **G.M.** Data modelling. **Y.W.** Image analysis. **Y.P.Z.**, **J.Y.L.L.** Data collection. **F.W.**, **W.A.M.** Providing the mouse samples. **A.Q**. Providing human brain samples, neuropathological characterisation. **M.G.S.** Conception and design, providing the mouse samples, manuscript preparation. **J.S.H.D.** Conception and design, data collection and image analysis. **D.K.** Conception and design, manuscript preparation, overall supervision of the project.

## Acknowledgments

We thank Dr. Marc Aurel Busche for providing valuable feedback on the manuscript. EM specimen preparation and TEM were performed using the facilities at CAIC (Cambridge Advanced Imaging Centre). We thank Karin Müller, Filomena Gallo, and Georgina Lindop for their kind support and help.

## 7. Figure captions

**Supplemental Figure 1.** Co-localisation of bassoon (pre-synaptic marker), PSD95 (post-synaptic marker), and CellMask confirms the successful isolation and localisation of synaptosomes on the SiMPull surface.

