## Supplementary figures and images for "SynPull: a novel method for studying neurodegeneration-related aggregates in synaptosomes using super-resolution microscopy"

### Supplemental Figure 1

Supp. Fig. 1

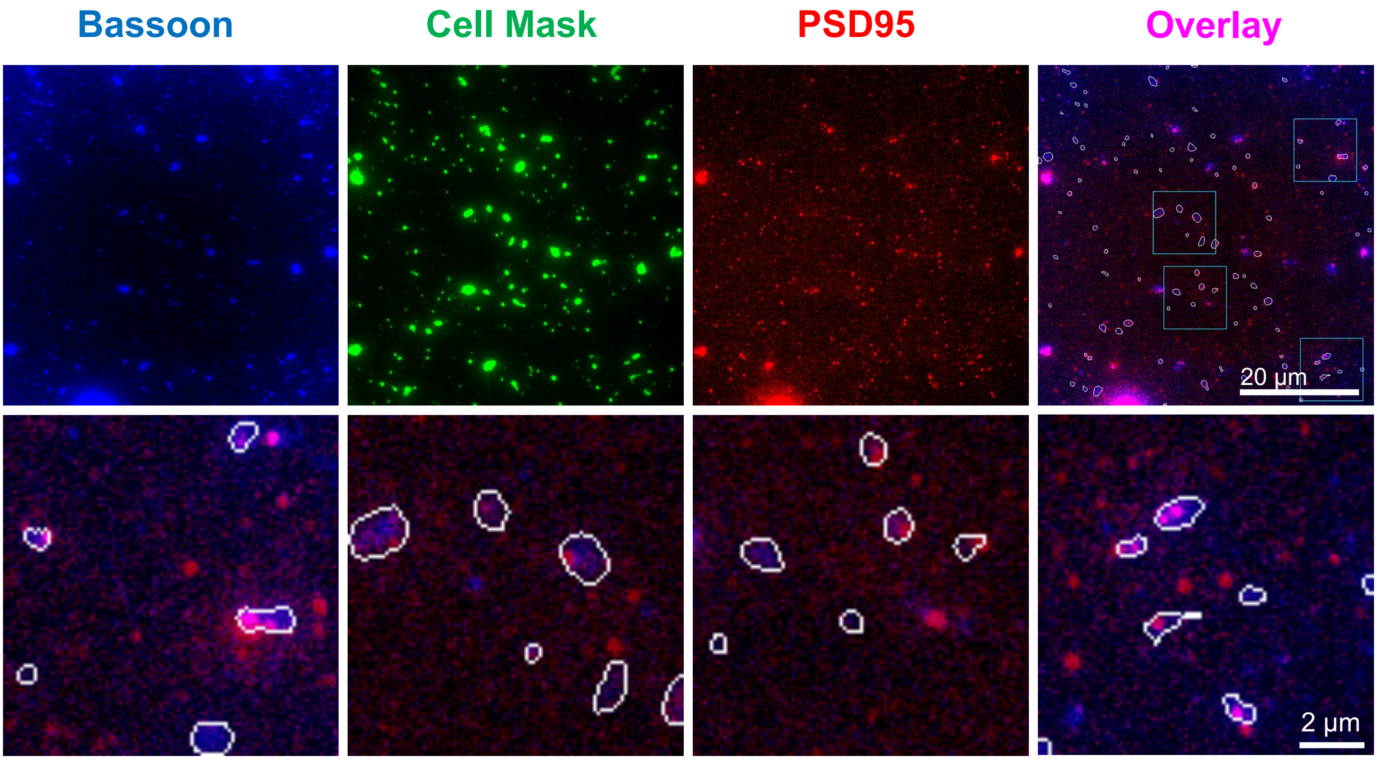
